# Engineered kinesin motor proteins amenable to small molecule inhibition

**DOI:** 10.1101/042663

**Authors:** Martin F. Engelke, Michael Winding, Yang Yue, Shankar Shastry, Federico Teloni, Sanjay Reddy, T. Lynne Blasius, Pushpanjali Soppina, William O. Hancock, Vladimir I. Gelfand, Kristen J. Verhey

**Affiliations:** Department of Cell and Developmental Biology, University of Michigan, Ann Arbor, MI 48109; Department of Cell and Molecular Biology, Northwestern University Feinberg School of Medicine, Chicago, IL 60611; Department of Biomedical Engineering, Penn State University, University Park, PA 16802; Current Address: Department of Chemistry and Biochemistry, UC Santa Cruz, Santa Cruz, CA 95064

## Abstract

The human genome encodes 45 kinesins that drive cell division, cell motility, intracellular trafficking, and ciliary function. Determining the cellular function of each kinesin would be greatly facilitated by specific small molecule inhibitors, but screens have yielded inhibitors that are specific to only a small number of kinesins, likely due to the high conservation of the kinesin motor domain across the superfamily. Here we present a chemical-genetic approach to engineer kinesin motors that retain microtubule-dependent motility in the absence of inhibitor yet can be efficiently inhibited by small, cell-permeable molecules. Using kinesin-1 as a prototype, we tested two independent strategies to design inhibitable motors. First, we inserted the six amino acid tetracysteine tag into surface loops of the motor domain such that binding of biarsenic dyes allosterically inhibits processive motility. Second, we fused DmrB dimerization domains to the motor heads such that addition of B/B homodimerizer cross-links the two motor domains and inhibits motor stepping. We show, using cellular assays that the engineered kinesin-1 motors are able to transport artificial and natural kinesin-1 cargoes, but are efficiently inhibited by the addition of the relevant small molecule. Single-molecule imaging *in vitro* revealed that inhibitor addition reduces the number of processively moving motors on the microtubule, with minor effects on motor run length and velocity. It is likely that these inhibition strategies can be successfully applied to other members of the kinesin superfamily due to the high conservation of the kinesin motor domain. The described engineered motors will be of great utility to dynamically and specifically study kinesin function in cells and animals.

## INTRODUCTION

Microtubules and associated proteins drive a variety of cellular processes including cell division, cell motility, intracellular trafficking, and ciliary function. Two motor protein families, kinesins and dyneins, produce force and/or motility along microtubule polymers and defects in these motors are associated with human pathologies including neurodegeneration, tumorigenesis, developmental defects, and ciliopathies^1-4^. Kinesins contain a highly conserved ~350 amino acid kinesin motor domain with signature sequences for ATP hydrolysis and microtubule binding. Many kinesins undergo processive motility and advance along the microtubule surface as dimeric molecules by alternate stepping of the two motor domains^5^. Outside of the motor domain, each kinesin contains unique sequences for cargo binding and regulation, and thereby carries out specific cellular transport functions^6, 7^.

Mammals contain ~ 45 kinesin genes that are classified into 17 families based on phylogentic analysis^8^. To identify the cellular roles of specific kinesin gene products, genetic approaches (e.g. knockout animals) and classical protein inhibition methods (e.g. RNAi, overexpression of dominant-negative proteins, injection of inhibitory antibodies) have been utilized. However, these approaches are hampered by off-target effects, gradual inhibition of the targeted kinesin, and/or the lack of timely control over protein inhibition and are thus not optimal for dissecting complex and dynamic cellular pathways. These drawbacks could in principle be overcome by the use of cell-permeable inhibitors, but screening efforts with small molecule libraries have yielded only few specific inhibitors^9^; most inhibitors target multiple kinesin motors, presumably due to the high conservation of the kinesin motor domain^10, 11^.

Here we report an “engineered chemical-genetic” approach to generate kinesin motors that are amenable to small molecule inhibition. Using kinesin-1 as a prototype, we developed two independent strategies to engineer genetically-modified motors that transport cellular cargoes in a manner indistinguishable from the wild-type motor but that can be rapidly and specifically inhibited with high specificity by the addition of a small molecule. Our approach enables investigation of the function of the kinesin-1 motor protein in cells or animals with high temporal resolution and specificity. Furthermore, due to the high conservation of the motor domain across the kinesin superfamily, our approach can likely be generalized to generate inhibitable versions of any kinesin motor of interest.

## RESULTS

### Designing kinesin motors amenable to small-molecule inhibition

Kinesins that are engineered to serve as tools to study motor function in cells and animals must fulfill two criteria. First, the engineered motor must maintain the microtubule-dependent motility properties of the wild-type protein and second, it must be specifically inhibited by a small, membrane-permeable molecule. Thus, a successful design will minimally alter the structure of the motor yet will mediate binding of the inhibitory molecule with high specificity and affinity. We pursued two strategies to yield kinesins that can be inhibited by addition of a small molecule. Both strategies were implemented and tested with kinesin-1 because it is the best-characterized member of the kinesin family and assays to study its motility and function are well-established [e.g. ref. 12-19].

Our first strategy to engineer inhibitable kinesin-1 motors took advantage of the ability of membrane-permeable biarsenical dyes (FlAsH and ReAsH) to bind to the small tetracysteine tag (TC, amino acid sequence CCPGCC) and thereby label TC-tagged proteins in live cells^20, 21^. We hypothesized that when the TC tag is inserted into the surface of the kinesin motor domain it will, in a ligand-dependent manner, restrict the conformational changes that occur during the catalytic cycle and thereby inhibit the motor (**Fig. 1a**). This strategy was first tested in *D. melanogaster* kinesin heavy chain (*Dm*KHC), where the TC tags were introduced into six different loops (L1, L2, L3, L5, L8, L14) on the surface of the motor domain in the active motor *Dm*KHC(1-559) (**Fig. 1c**, **Supplementary Fig. 1a-c**, and ref. 22). Microtubule gliding assays with purified motors revealed that insertion of the TC tag into L3, L5 or L8 abolished motor activity (**Table 1**) and these constructs were not pursued further. Insertion of the TC tag into L14 resulted in motors with reduced gliding speed that was not altered by the presence of the FlAsH dye (**Table 1**) and this construct was also not further pursued. Importantly, insertion of the TC tag into L1 or L2 resulted in motors that had gliding activity similar WT motor in the absence of inhibitor and no measurable gliding activity in the presence of the FlAsH dye (**Table 1**). We thus pursued the insertion of a TC motif into L1 or L2 for generating an inhibitable mammalian kinesin-1 motor using a truncated and constitutively active version of the *R. norvegicus* kinesin-1 motor *Rn*KIF5C(1-559) (**Fig. 1b,c** and **Supplementary Fig. 2a,b**).

**Table 1.**
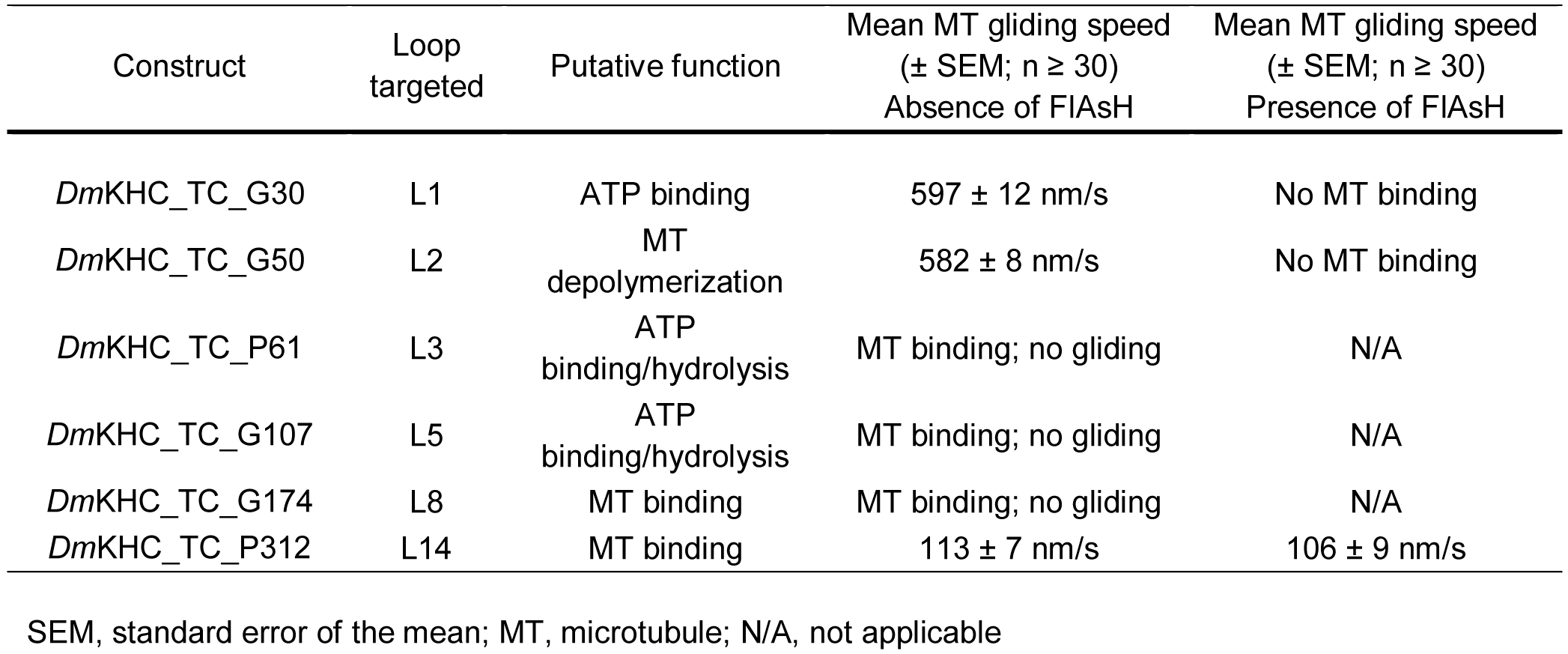
Microtubule gliding assay with TC-engineered *Dm*KHC constructs

SEM, standard error of the mean; MT, microtubule; N/A, not applicable

**Figure 1.**
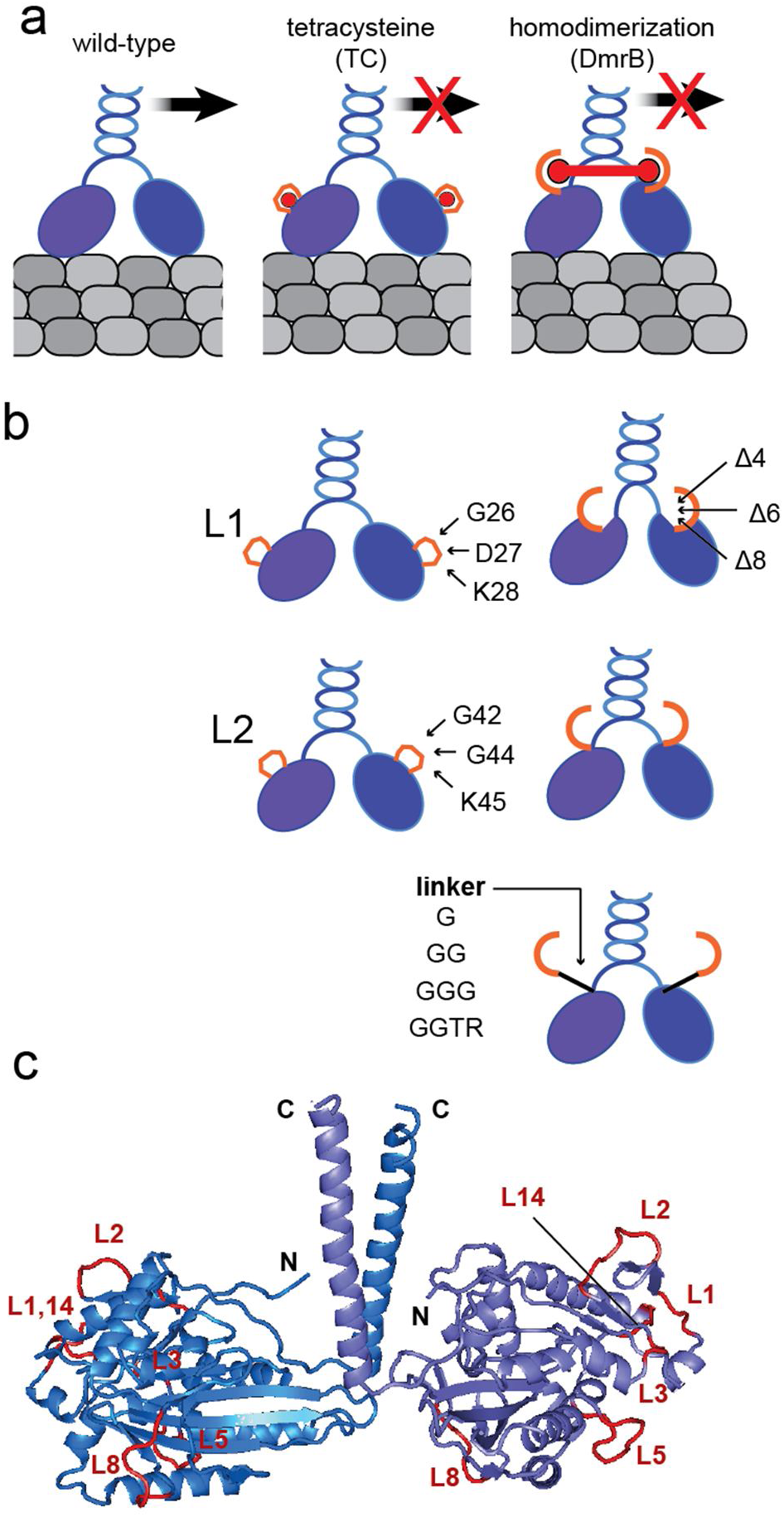
Strategies to make kinesin-1 responsive to small molecule inhibition. (**a**) The two motor domains (blue ovals) of a dimeric kinesin-1 undergo alternating catalysis to generate processive motility along the microtubule surface (gray). The tetracysteine (TC) strategy inserts a TC tag (orange loop; sequence CCPGCC) into surface loops of the kinesin motor domain. Binding of the biarsenic dye ReAsH (red circle) to the TC tag restricts the conformational changes of the motor domain during catalysis, resulting in allosteric inhibition of microtubule-dependent motor motility. The homodimerization (DmrB) strategy fuses a DmrB domain (orange semicircle) to the N-terminus of the kinesin motor domain. Addition of the B/B homodimerizer (red dumbbell) to the DmrB domains cross-links the motor domains and inhibits processive motility. (**b**) Schematic showing the engineered TC and DmrB constructs. TC tags were incorporated at the indicated positions in L1 (G26,D27,K28) or L2 (G42,G44,K45) of a truncated *Rn*KIF5C(1-559) motor. The DmrB domain was fused directly to the N-terminus (middle), extended by short peptide linkers (G, GG, GGG, or GGTR, bottom) or brought closer by deletion of the N-terminal 4, 6, or 8 amino acids of kinesin-1 (A4, A6, A8, top). (**c**) Ribbon diagram of the rat kinesin-1 dimeric motor domain (PDB 3KIN). The TC tag was inserted into L1 and L2 on the surface of the motor domain to generate engineered *Rn*KIF5C(1-559) and into L1, L2, L3, L5, L8, and L14 to generate the *Dm*KHC(1-559) constructs, whereas DmrB was fused to the N-terminus (N). C, C-terminus

Our second strategy to engineer inhibitable kinesin-1 motors was based on the ability of the cell-permeable rapamycin analog B/B homodimerizer (Rapalog-2 or AP20187) to induce homodimerization of the DmrB domain (F36V variant of FKBP^23, 24^). We reasoned that fusion of the DmrB domain to the N-terminus of the kinesin motor domain will create a situation in which addition of B/B homodimerizer (hereafter called B/B) cross-links the motor domains of the kinesin dimer and inhibits motor stepping (**Fig. 1a**). We created a series of constructs in which the DmrB domain was fused directly to the N-terminus of *Rn*KIF5C(1-559), separated from the N-terminal methionine by one (DmrB-G), two (DmrB-GG), three (DmrB-GGG), or four (DmrB-GGTR) amino acids, or situated closer to the core motor domain by removing the N-terminal four (DmrBΔ4), six (DmrBΔ6) or eight (DmrBΔ8) amino acids of KIF5C (**Fig. 1b,c**).

### Initial analysis of engineered mammalian kinesin-1 motors

We first verified that the engineered TC and DmrB *Rn*KIF5C(1-559) motor dimers retain microtubule-dependent motility in the absence of the inhibitors. To do this, we expressed the constructs in differentiated CAD cells where the defined microtubule architecture of the neuronal processes enables the accumulation of active kinesin motors at the distal tip^25^. All TC constructs showed accumulation in neurite tips (**Supplementary Fig. 3**), suggesting that insertion of the TC motif into L1 or L2 does not interfere with microtubule-dependent motility. For the DmrB constructs, fusion of the DmrB domain directly to the N-terminus or separating it from the N-terminus via short linkers resulted in motor accumulation at neurite tips (**Supplementary Fig. 3**). However, positioning the DmrB domain closer to the core motor domain reduced or abolished tip accumulation (**Supplementary Fig. 3**), consistent with reports that these N-terminal residues form a cover strand essential for motor force generation^26, 27^. The DmrBΔ4, DmrBΔ6, and DmrBΔ8 constructs were thus excluded from further consideration. TC_G44 was also not considered further as this construct appeared to aggregate in some cells (data not shown). All remaining TC and DmrB constructs generate stable proteins with the expected molecular weights (**Supplementary Fig. 2c**), and were thus pursued in further detail.

To analyze motor activity in the absence and presence of the inhibitors, we performed single-molecule imaging of 3xmCit-tagged motors using total internal reflection fluorescence (TIRF) microscopy. (**Fig. 2a**). We found that addition of ReAsH to TC-tagged KIF5C(1-559)-3xmCit motors or addition of B/B to DmrB-tagged KIF5C(1-559)-3xmCit motors resulted in a significant reduction of the motor landing rate, which is defined as the frequency with which motors land and processively move (> 250 nm) along the microtubule (**Fig. 2b** and **Supplementary Fig. 4a**). However, the run velocity and run length of the motors in the presence of the inhibitors were not significantly different than that of the WT motor (**Fig. 2c,d** and **Supplementary Fig. 4b,c**). These data indicate that the ReAsH and B/B inhibitors directly interfere with the ability of the engineered motors to engage with the microtubule track in a constructive manner.

**Figure 2.**
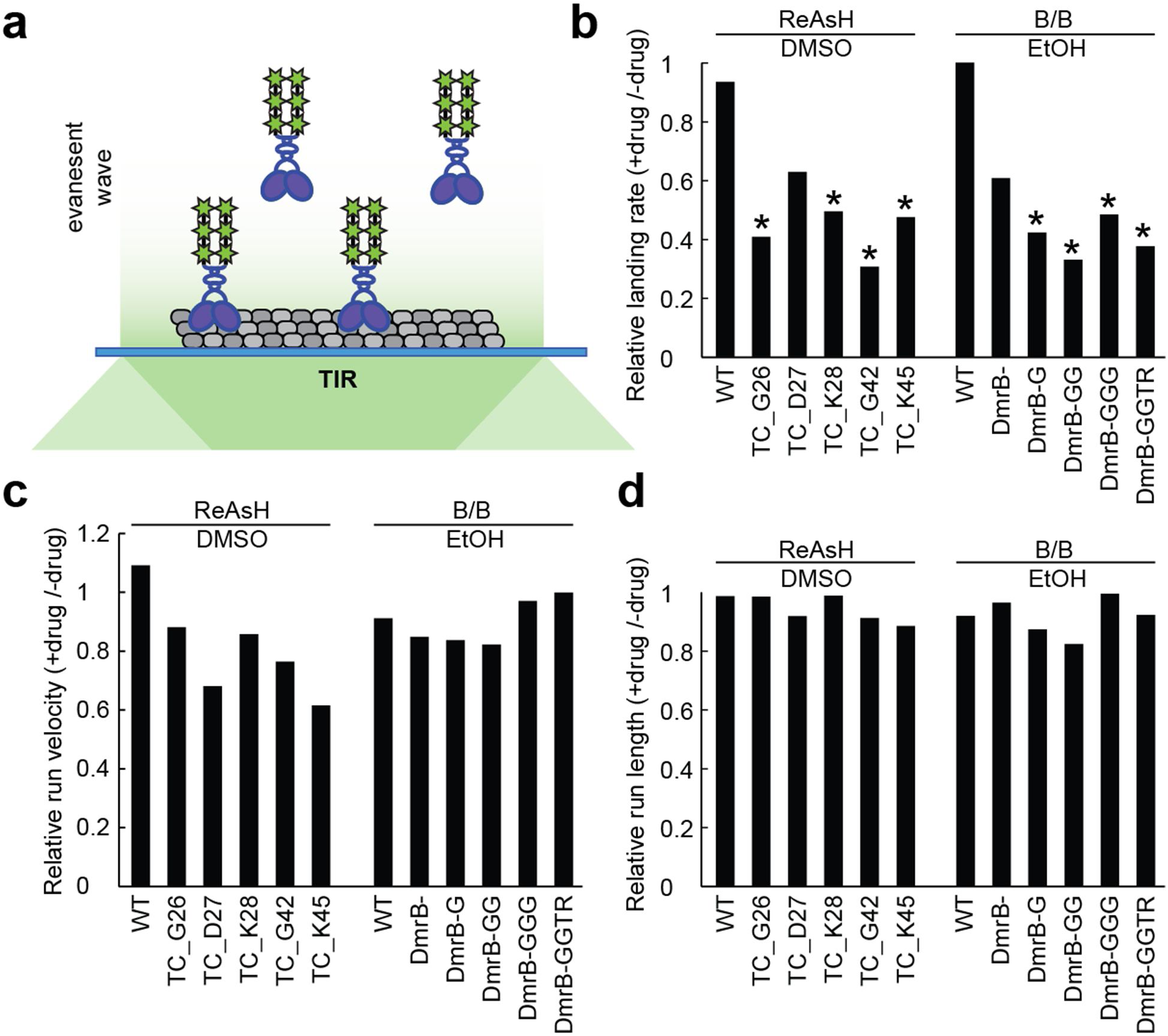
Effects of inhibition on the single molecule motility properties of engineered kinesin-1 motors. (**a**) Schematic of the *in vitro* assay for imaging single *Rn*KIF5C(1-559) motors tagged with tandem mCit fluorescent proteins (green stars). TIR, total internal reflection. (**b-d**) Lysates of COS7 cells expressing WT, TC-tagged or DmrB-tagged *Rn*KIF5C(1-559) motors were incubated for 30 min in the absence (EtOH, DMSO) or presence (B/B, ReAsH) of the respective inhibitor. The processive motility of single motors along taxol-stabilized microtubules was observed by TIRF microscopy. The landing rate, run velocity and run length were determined from 7-9 time-lapse movies from three independent experiments. The data are presented as the relative change in the mean (**b**) landing rate, (**c**) run velocity and (**d**) run length obtained +drug /-drug. * p<0.05; heteroscedastic, two-tailed t test comparing motility frequency in the absence and presence of drug.

### Rapid inhibition of engineered TC motors in cells

To characterize the inhibitable kinesin-1 motors in terms of cargo transport and inhibition in cells, we employed an inducible peroxisome redistribution assay in which recruitment of active kinesin motors to peroxisomes causes a redistribution of the peroxisomes from their perinuclear localization to the cell periphery^28^ (**Fig. 3a**) and hence provides a fast and direct readout of motor activity. For this assay, the DmrA (FKBP) domain is targeted to the peroxisome surface via a PEX sequence^29^ and rapamycin or A/C heterodimerizer (Rapalog-1 or AP21967) are used to recruit DmrC (FRB)-tagged kinesin motors to peroxisomes in a rapid manner. Unfortunately, this assay could not be used to probe the DmrB constructs due to cross-reaction of the A/C heterodimerizer with the DmrB domain.

**Figure 3.**
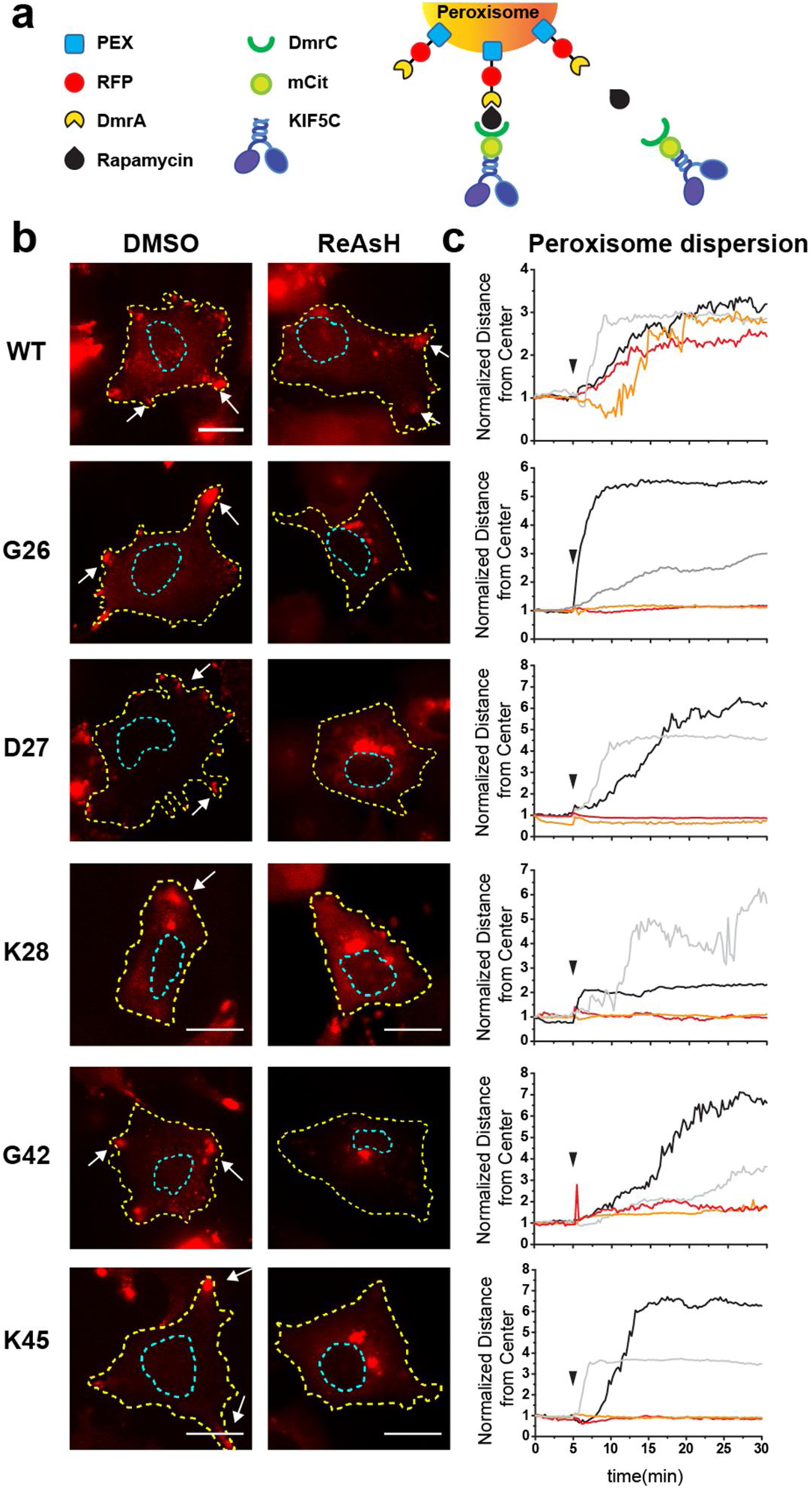
Rapid inhibition of TC-tagged kinesin-1-mediated transport in cells. (**a**) Schematic diagram of the inducible peroxisome transport assay. WT-or TC-tagged *Rn*KIF5C(1-559) motor constructs fused to mCitrine (mCit) and a DmrC domain can be recruited to the surface of peroxisomes containing a PEX-RFP-DmrA construct by addition of A/C heterodimerizer (or rapamycin). Recruitment of active motors results in dispersion of peroxisomes to the cell periphery. (**b,c**) Cells expressing the indicated WT or TC-tagged motor constructs were treated with DMSO (vehicle control) or ReAsH for 30 min. Cells were then imaged for 30 min and rapamycin was added 5 min after the start of imaging to induce motor recruitment to the peroxisome surface. (**b**) Representative images of the peroxisome distribution at 30 min. Yellow dotted line, cell periphery; blue dotted line, nucleus. Arrows indicate the accumulation of peroxisomes in the periphery of the cell. Scale bar, 20 μm. (**c**) Time course of peroxisome redistribution in DMSO (black and light grey lines) or ReAsH (red and orange lines) treated cells. The arrow head at 5 min indicates the time of rapamycin addition. The data are presented as the average distance of the peroxisomes normalized to the average starting positions in each cell.

Recruitment of WT or TC-tagged KIF5C(1-559) motors to the peroxisome surface in control treated cells (DMSO) resulted in a redistribution of the peroxisomes to the cell periphery, as expected (**Fig. 3b,c** and **Supplementary Fig. 5**). Addition of ReAsH dye had no effect on peroxisome redistribution driven by WT motors but abolished the redistribution driven by TC motors (**Fig. 3b,c** and **Supplementary Fig. 5**). These results demonstrate that addition of ReAsH efficiently inhibits the transport function of engineered TC motors within 30 min of inhibitor addition. We also found that FlAsH dye is less potent than ReAsH in blocking peroxisome distribution by engineered TC motors (data not shown). It is possible that the more bulky fluorescein moiety of FlAsH sterically hinders its binding to the TC tag on the surface of the kinesin motor domain.

### Engineered motors transport artificial cargo in the absence but not presence of inhibitor

To further examine the ability of the engineered TC and DmrB motors to be inhibited by the relevant small molecule in cells, we utilized a Golgi redistribution assay^30^ in which Golgi-targeted active kinesin-1 motors cause a redistribution of the Golgi complex from its characteristic tightly-packed perinuclear location to a dispersed phenotype of small Golgi-derived vesicles scattered throughout the cytoplasm (**Fig. 4a,b**). This assay shows less cell-to-cell variability (data not shown) and avoids the cross-reaction of the DmrA/C heterodimerizer with the DmrB domain in the inducible peroxisome redistribution assay.

**Figure 4.**
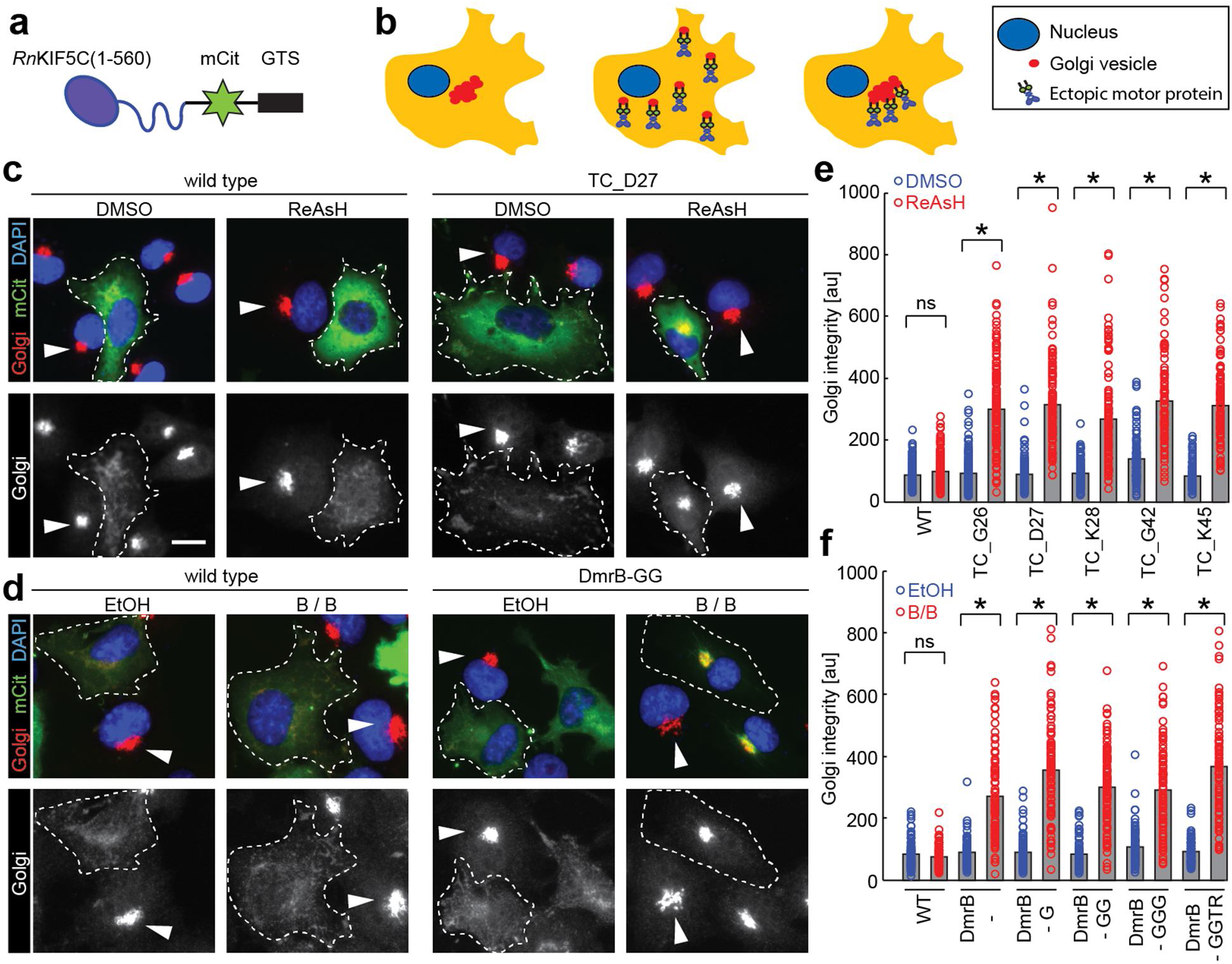
Inhibition of transport of artificial cargo by engineered kinesin-1 motors in cells. (**a**) Schematic of *Rn*KIF5C(1-559) motor tagged with mCit and a Golgi targeting sequence (GTS). (**b**) Schematic of Golgi redistribution assay. In untransfected cells (left), the Golgi complex (red) is localized in a tightly-packed, perinuclear cluster. Expression of active kinesin-GTS motors (middle) results in redistribution of the Golgi complex to a dispersed localization whereas inhibition of these motors (right) retains the tightly-packed, perinuclear Golgi localization. (**c-f**) COS7 cells expressing WT, TC-tagged or DmrB-tagged motors with a GTS were incubated in the absence (DMSO or EtOH) or presence of inhibitor (200-400 nM ReAsH or 1.5 μM B/B), fixed ~14h later, and then stained with DAPI (blue) and an antibody to a Golgi marker (Giantin; red). The top panels show merged images whereas the bottom panels show only the Golgi fluorescence signal. White dotted line, outline of cells expressing engineered motors. Arrowheads indicate compact Golgi complex in non-expressing cells. Scale bar, 15 μm. (**c**) Representative images from cells expressing WT or TC_D27 constructs. (**d**) Representative images from cells expressing WT or DmrB-GG constructs. (**e,f**) Quantification of Golgi redistribution. The data are presented as the Golgi integrity where low values reflect a dispersed Golgi complex (motors active) and high values reflect a compact perinuclear Golgi complex (motors inhibited). Each circle represents the Golgi integrity of an individual cell in the presence (red) or absence (blue) of the respective inhibitor. Gray bars represent means (**e**, n ≥ 100; **f**, n y 76) of three independent experiments. p-values were calculated with a heteroscedastic, two-tailed t test; ns, not significant; * p<0.000001

WT and engineered *Rn*KIF5C(1-559) motors were targeted to the Golgi complex by fusion of the Golgi targeting sequence (GTS) of the cis-Golgi resident protein GMAP210^30-32^. Golgi-targeting of WT, TC-tagged, and DmrB-tagged KIF5C(1-559) motors in the absence of any inhibitor resulted in a dispersed Golgi phenotype (**Fig. 4c-f**), suggesting that the engineered TC and DmrB motors are not only capable of processive motility but also generation of sufficient force to oppose the Golgi-localized dynein that is responsible for its perinuclear clustering. Addition of ReAsH dye blocked the Golgi dispersion driven by TC-tagged motors but not WT motors (**Fig. 4c,e**). Likewise, addition of B/B blocked Golgi dispersion driven by DmrB-tagged but not WT kinesin-1 motors (**Fig. 4d,f**). We found that in this assay, TC_G42 motors often formed small foci in the cytoplasm (**Supplementary Fig. 6**), suggesting that this motor may be prone to aggregation when overexpressed. Furthermore we observed that inhibition of the DmrB constructs was most efficient in cells expressing low levels of the engineered motor construct (based on mCit fluorescence, data not shown). While a concentration of 1.5 μM B/B homodimerizer was chosen based on previous work^33^, we found the optimal concentration of ReAsH inhibitor to be 200-400 nM (**Supplementary Fig. 7**).

To quantify this striking difference in Golgi morphology, we measured the standard deviation of pixel intensities for the Golgi staining in the perinuclear region on a cell-by-cell basis (**Online Methods**). Under control conditions (untransfected) or when the Golgi-targeted TC and DmrB motors are inhibited, the compact Golgi complex results in a high standard deviation of pixel intensities in the perinuclear region whereas the dispersed Golgi complex in cells expressing active WT or engineered motors displays a small standard deviation of pixel intensities. Since this quantification method measures pixel intensity fluctuations around the mean in a subcellular region, it is not sensitive to different staining efficiencies and exposure times and is hence suited to comparing conditions across independent experiments. Quantification of the Golgi redistribution revealed that addition of the ReAsH or B/B inhibitors resulted in a significant inhibition of Golgi transport by the TC and DmrB constructs, respectively (**Fig. 4e,f**). These results demonstrate that the TC and DmrB kinesin-1 constructs are capable of cargo transport in cells and that this transport can be specifically inhibited upon addition of the relevant small molecule inhibitor.

### Inhibitable motors rescue endogenous motor function and can be efficiently inhibited in cells

As a final test of the inhibitable kinesin-1 motors, we examined whether the engineered motors are able to rescue the function of the endogenous motor in cells and whether the transport of endogenous cargoes can be inhibited by the addition of the cognate inhibitor molecule. To do this, we chose *D. melanogaster* S2 cells because i) expression of the single *Drosophila* kinesin-1 gene, *Dm*KHC, can be efficiently knocked down using dsRNA against the 3’-UTR of the mRNA and ii) *Dm*KHC is solely responsible for microtubule-microtubule sliding in these cells, thus providing a direct readout of kinesin-1 activity^34^.

We created full-length versions of the inhibitable, TC-tagged *Dm*KHC constructs G30 (L1) and G50 (L2) (**Table 1** and **Supplementary Fig. 1a-c**). We also created new DmrB-tagged versions of full-length *Dm*KHC based on our success with the DmrB-tagged *Rn*KIF5C(1-559) motors. Since *Dm*KHC contains four additional amino acids on the N-terminus of its motor domain (**Supplementary Fig. 1d**), we fused the DmrB-domain directly to the N-terminus of *Dm*KHC (= DmrB-*Dm*KHC, analogous to the *Rn*KIF5C(1-559) construct DmrB-GGTR) or removed three amino acids from the DmKHC N-terminus (= DmrB-Δ3*Dm*KHC, analogous to the *Rn*KIF5C(1-559) construct DmrB-G). To monitor expression, a blue fluorescent protein tag, mTagBFP2 (BFP), was fused to the C-terminus of each construct.

Microtubule sliding was measured in S2 cells by expressing tubulin tagged with a tandem dimer of mEOS2 (tdEOS-tubulin)^18, 35^, photoconverting an area of green fluorescent microtubules to red fluorescence, and measuring the movement of the red fluorescent microtubules over time (**Fig. 5a,b** and **Supplementary Video 1**). Knockdown of endogenous *Dm*KHC significantly reduced the microtubule sliding rate in comparison to control treated cells, as reported previously^34^, and microtubule sliding was efficiently rescued by the ectopic expression of BFP-labeled WT, TC-tagged or DmrB-tagged motors (**Fig. 5c,d**). Importantly, addition of ReAsH dye blocked the activity of the *Dm*KHC TC_G30 and TC_G50 motors as microtubule sliding was reduced to that of the knockdown cells (**Fig. 5c**). Likewise, addition of B/B homodimerizer blocked the activity of the DmrB-*Dm*KHC and DmrB-Δ3*Dm*KHC motors (**Fig. 5d**). These results demonstrate that each of the engineered DmrB or TC *Dm*KHC constructs can rescue the activity of the wild-type motor and that engineered motors can be efficiently inhibited by the cognate inhibitors.

**Figure 5.**
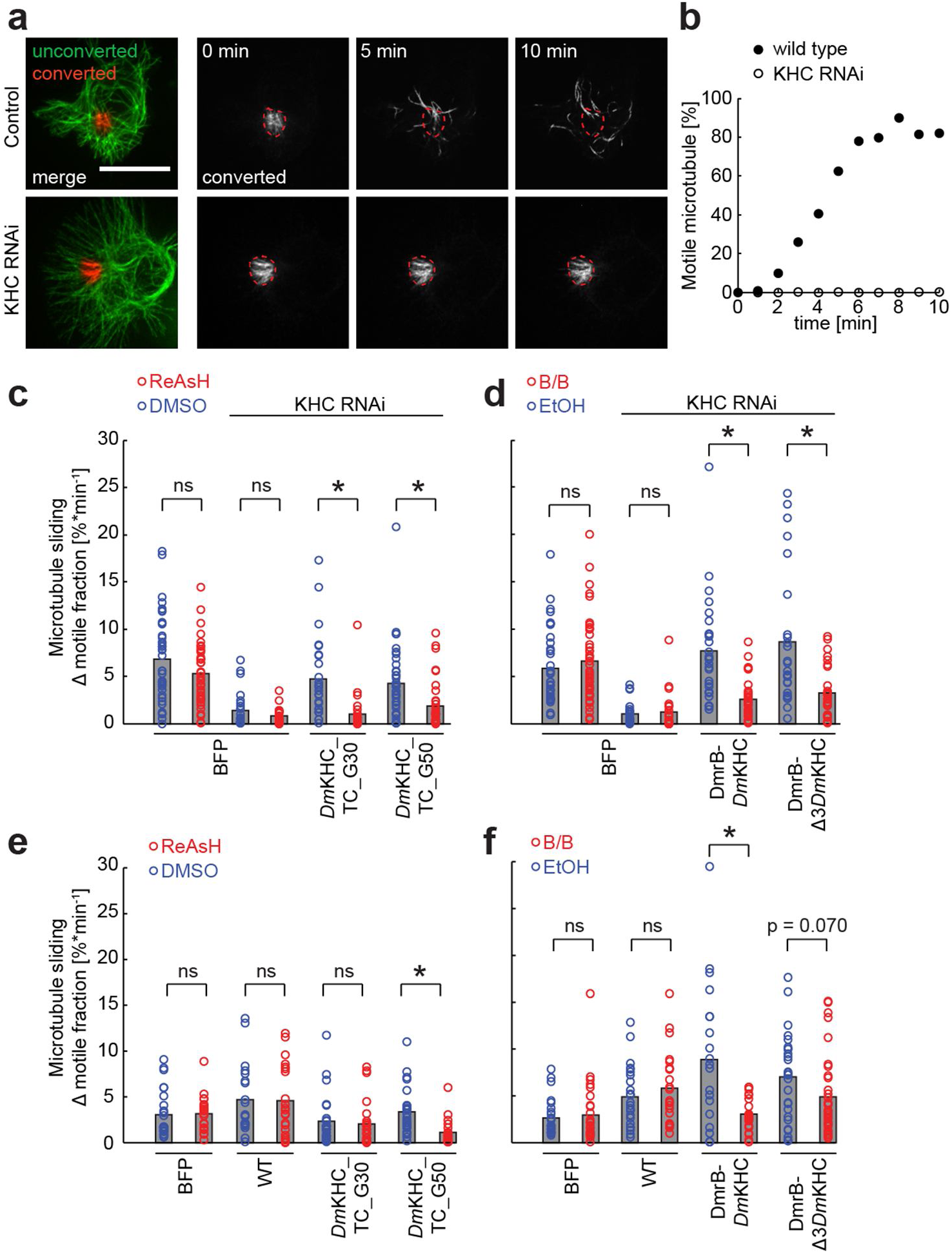
Inhibition of kinesin-1 function in *D. melanogaster* S2 cells expressing engineered *Dm*KHC motors. Microtubule sliding was measured in cells expressing tdEOS-tubulin under control (no RNAi), kinesin-1 knockdown (KHC RNAi), or upon knockdown and rescue with blue fluorescent protein (BFP) alone or BFP-tagged WT, TC, or DmrB *Dm*KHC motors. On the day of imaging, cells were treated with vehicle (DMSO or EtOH) or inhibitor (200-400 nM ReAsH or 1.5 μM B/B) and tdEOS-tagged microtubules were photoconverted from green to red fluorescence in a region of interest. Photoconverted microtubules were tracked over a period of 5-10 min to measure microtubule sliding, a measure of kinesin-1 activity. (**a**) Representative images of microtubule sliding over time in control (no RNAi) or KHC RNAi-treated S2 cells. Scale bar, 15 μm. (**b**) Quantification of the microtubule sliding rates from (**a**). (**c-f**) Microtubule sliding driven by engineered *Dm*KHC motors in (**c,d**) the absence of endogenous *Dm*KHC (KHC RNAi) or (**e,f**) the presence of endogenous *Dm*KHC. Each circle represents the microtubule sliding activity of a single cell. The mean of each condition (**c-d**: n ≥ 25; **e-f**: n ≥ 20) is derived from values of at least two independent experiments and is shown by a gray bar. High values reflect high *Dm*KHC activity whereas low values reflect low *Dm*KHC activity. p-values were calculated with a heteroscedastic, two-tailed t test; ns, not significant; * p<0.005.

We then used the microtubule-microtubule sliding in S2 cells as an assay to test whether the engineered kinesin-1 motors can inhibit transport in the presence of the endogenous motor. Expression of *Dm*KHC TC_G30 surprisingly had a weak dominant-negative effect on endogenous kinesin-1 activity, but this effect is not significant and microtubule sliding was not further reduced by addition of ReAsH (**Fig. 5e**). In contrast, addition of ReAsH reduced microtubule sliding in cells expressing TC_G50 (**Fig. 5e**), suggesting that kinesin motors containing an insertion of TC in L2 can inhibit the function of the endogenous motor in trans. For the DmrB-tagged motors, addition of B/B reduced the microtubule sliding rate in cells expressing DmrB-*Dm*KHC and DmrB-Δ3*Dm*KHC but not below the level driven by the endogenous motor (**Fig. 5f**), suggesting that when inhibited, the DmrB motors do not interfere with the function of the endogenous motor.

## DISCUSSION

Chemical-genetic approaches have been successfully used to study the function of proteins for which the identification of specific inhibitors has remained elusive^36, 37^. For ATPases such as kinases or motor proteins (including myosins and kinesins), such approaches have involved enlargement of the ATP binding pocket to accommodate a nucleotide or inhibitor analogue that contains a bulky substituent complementing the enlarged pocket. For some kinases, modulation of the gatekeeper residue resulted in diminished kinase activity and ATP affinity and recent efforts have focused on addressing this challenge^38^. Analog-sensitive versions of kinesin-1 have been generated but have not been used for cell biological studies as the engineered motor requires a modified ATP for full motor activity and the modified nucleotides are not membrane permeable^39^.

Here we describe a novel chemical-genetic approach to inhibit kinesin-1 that has several advantages over existing inhibition method. First, our approach is based on cell-permeable inhibitors and is thus minimally invasive and tunable. Second, we use small molecules that bind their cognate target peptide/domain with high affinity and specificity, granting fast and selective inhibition of the targeted kinesin. Third, the engineered kinesins have properties similar to wild type kinesins, making the system ideal for acute inhibition in cellular/developmental systems. Fourth, the DmrB strategy is based on the established B/B homodimerizer (Rapalog-2), a chemical derivative of rapamycin with well-studied pharmacokinetics and the possibility of photocaging for enhanced spatial control in cellular systems^40, 41^.

We demonstrate that DmrB-or TC-tagging of rat and fly kinesin-1 results in motor constructs that display similar microtubule-dependent motility properties as wild-type motors but can be inhibited with small, cell-permeable molecules in three types of assays: as single-molecules *in vitro*, driving transport of exogenous cargoes in cells, and driving transport of an endogenous cargo in cells. It is interesting to note that the level of inhibition was greater for cargo transport assays in cells than for single motors *in vitro*, perhaps due to an incomplete inhibition of motors *in vitro* and/or the requirement for inhibitor-bound motors to produce force in cells but not *in vitro*. For the rat KIF5C motor, placement of the TC tag in six different locations in L1 or L2 resulted in inhibition although differences were observed between constructs in terms of protein expression and/or aggregation. For future work, we favor the constructs *Dm*KHC_TC_G30 and the corresponding rat TC_D27 since these constructs consistently performed well in all assays. In addition, L1 is more conserved across kinesin family members than L2, making the transfer of this approach to other kinesin motors easier to implement for L1. For the DmrB-tagged rat and fly kinesin-1 motors, we slightly favor the rat construct DmrB-GG (corresponding to the fly construct DmrB-*Dm*KHC), because it shows the highest degree of inhibition at the single-molecule level (**Fig. 2b**) and a strong effects in the Golgi dispersion assay (**Fig. 4f**). The N-terminal residues vary widely between kinesins and the placement of the DmrB domain will depend on the length of the N-terminal extension. In order to fully utilize the inhibitable kinesin-1 constructs, the engineered constructs should be expressed in cells in which endogenous kinesin-1 is either absent or has been depleted (**Fig. 5c-f**).

In summary, the kinesin inhibition approach that we describe enables the specific and dynamic perturbation of kinesin function. This approach will enable researchers to decipher the exact roles of kinesin-1 in various transport events in specific cellular/developmental stages and facilitate our understanding of the interactions between motors during a tug-of-war. We anticipate that our approach will be applicable in animals where the endogenous kinesin-1 has been genetically replaced with our engineered constructs. Given the high conservation of the kinesin motor domain across the kinesin superfamily, we also anticipate this approach will be directly transferable to other kinesin family members.

## ONLINE METHODS

### Plasmids and production of dsRNA

Sequences encoding tetracysteine tags (CCPGCC) were introduced into a *Dm*KHC truncated at position 559 and containing a C-terminal eGFP and His6 tags using Agilent Technologies? QuickChange XL II kit or by using Splice by Overlap Extension (SOE) PCR^42, 43^ to generate plasmids encoding constructs used in *in vitro* microtubule gliding assays. Sequences encoding TC-or DmrB-tagged motors were synthesized (DNA2.0 or Life Technologies) and cloned in-frame into *Rn*KIF5C(1-559)-3xmCit^44^. For the peroxisome targeting assay, the last two mCits of the WT or TC-tagged motors were replaced by a DmrC domain (Clontech). The peroxisome anchor construct PEX3-mRFP-2xFKBP was a gift from Dr. C.C. Hoogenraad (Utrecht University, Netherlands; ref. 45). For the Golgi redistribution assay, the Golgi-targeting sequence of *Hs*GMAP210 (amino acids 1757-1838, NP_004230) was synthesized (Life Technologies) and subcloned in-frame at the C-terminus of the WT or engineered motors. For microtubule sliding assays in S2 cells, sequences encoding TC-or DmrB-tagged motors were synthesized (Life Technologies) and subcloned in-frame into the pMT/V5-His A vector (Invitrogen) containing full-length *Dm*KHC with an mTagBFP2 fluorescent protein at the C-terminus. KIF16B-mCit has been described^46^. All constructs were verified by analytical restriction digestion and sequencing.

Double-stranded RNA was generated by *in vitro* transcribing RNA strands with T7 polymerase followed by LiCl purification. Template DNA was generated by PCR of genomic DNA obtained from white-eyed (w1118) adult flies. The 5’ end of each primer used in PCR reactions contained the T7 promoter sequence (TAATACGACTCACTATAGGG). The annealing sequence of the forward KHC 3′-UTR primer is ATCCAATCACCACCTGTCGC and the sequence of the reverse is TCTGCGACTTTTATTTAGGT.

### Cell culture, transfection, lysis, and immunofluorescence

COS7 cells (African green monkey kidney fibroblasts, ATCC) were cultured in D-MEM (Gibco) and CAD cells (mouse catecholaminergic cell line) in D-MEM/F12 (Gibco) medium supplemented with 10% Fetal Clone III (HyClone) and GlutaMAX (Gibco) and grown at 37°C and 5% CO_2_. *D. melanogaster* S2 cells were maintained in serum-free Insect-Xpress media with L-glutamine (Lonza). COS7 cells were transfected with Lipofectamine 2000 (Life Technologies), CAD with Trans-It LT1 (Mirus Bio LLC), and S2 cells with Effectene Transfection Reagent (Qiagen), each according to the manufacturer’s instructions.

COS7 cells expressing the fluorescently-tagged motor constructs of interest were lysed 18 h post-transfection. Cells were trypsinized, harvested by low speed centrifugation, washed once in full culture medium and lysed in chilled Lysis Buffer (25 mM HEPES/KOH pH 7.4, 115 mM potassium acetate, 5 mM sodium acetate, 5 mM MgCl_2_, 0.5 mM EGTA, and 1% Triton X-100,) supplemented with 1 mM PMSF and protease inhibitor cocktail (P8340, Sigma) and 1 mM ATP. The lysate was cleared by centrifugation at 16.000g, 4°C for 10 min and aliquots were flash-frozen in liquid nitrogen and stored at −80°C. The amount of motor proteins across lysates was normalized by dot blot analysis on BioTraceNT nitrocellulose membranes using monoclonal antibodies to GFP (A6455; Invitrogen) and kinesin-1 (MAB1614, Covance). ImageJ (NIH) was used for densitometric analysis.

For immunofluorescence, cells were fixed with 4% (vol/vol) formaldehyde in PBS, quenched with 50 mM NH_4_Cl in PBS, permeabilized with 0.2% Triton X-100 in PBS, and blocked with blocking buffer (0.2% fish skin gelatin in PBS). Primary and secondary antibodies were applied in blocking buffer for 1 h at room temperature in the dark. Commercial antibodies used: polyclonal cis-Golgi marker giantin (PRB-114C, Covance), β-tubulin (clone E7; Developmental Studies Hybridoma Bank). Nuclei were stained with 4’, 6-diamidino-2-phenylindole (DAPI) and cover glasses were mounted in ProlongGold (Invitrogen). Images were collected on an inverted epifluorescence microscope (Nikon TE2000E) with a 40x 0.75 NA objective and Photometrics CoolSnapHQ camera and analyzed using ImageJ (NIH).

### CAD cell tip accumulation assay

CAD cells were seeded on cover glasses in growth medium, which was exchanged for serum-free medium 4 h later, followed by transfection. Two days later cells were fixed and stained. Motor fluorescence at the proximal and distal ends of the neurites was measured and the tip accumulation is reported as the ratio of the distal (tip) fluorescence to the proximal (shaft) fluorescence.

### Golgi dispersion assay

COS7 cells were seeded on cover glasses and treated 4 h later with inhibitor (ReAsH-EDT_2_, Molecular Probes or B/B homodimerizer, Clontech) or its solvent (DMSO, Sigma or ethanol, Decon Laboratories, Inc.) directly followed by the addition of transfection complexes. After 14 h, B/B-treated cells were fixed whereas ReAsH-treated cells were first washed three times for 1 min each in BAL wash buffer (Molecular Probes) at room temperature and then fixed. For quantitative analysis of Golgi distribution, a MATLAB (Mathworks) script was developed to determine the standard deviation of pixel intensities in the Golgi channel over a dilated nuclear area. A region of interest (ROI) representing the nucleus was manually selected and then extended outward for 10 pixels using a threshold-based method^47^. A high standard deviation of pixel intensities in the extended ROI is obtained for a tightly packed, perinuclear Golgi whereas a low standard deviation is obtained for cells in which the Golgi-derived fluorescence signal is spread throughout the cell.

### Inducible peroxisome redistribution assay

COS7 cells were plated onto glass-bottomed dishes (MatTek Corporation) and co-transfected 3 h later with plasmids coding for DmrC-tagged motors and the peroxisome anchor PEX3-mRFP-2xDmrA. Sixteen hours later, cells were washed with Opti-MEM (Gibco) and incubated for 30 min at 37°C and 5% CO_2_ in Opti-MEM containing 0.1% DMSO (solvent control) or 200 nM ReAsH-EDT_2_ (Molecular Probes). Cells were then washed at room temperature three times for 1 min each with BAL wash buffer (Molecular Probes) and immersed in Leibovitz’s L-15 medium (Gibco) supplemented with 200 μM Trolox (Sigma-Aldrich). Live cell imaging was carried out at 37°C in a temperature-controlled and humidified live imaging chamber (Tokai Hit) on a Nikon Ti-E/B microscope equipped with a 100x 1.49 NA oil immersion TIRF objective, three 20 mW diode lasers (488 nm, 561 nm, 640 nm), and EMCCD detector (iXon X3 DU897, Andor). Images were recorded in both the 488 nm and 561 nm channels every 15 s for 5 min and then 40 ng/ml rapamycin (Calbiochem) was added to the imaging medium and imaging resumed for another 25 min. Shifts in cell position during imaging were corrected with the Template Matching and Slice Alignment plugin to ImageJ (NIH) and time-lapse images were analyzed with a custom MATLAB (Mathworks) script. First peroxisome objects were detected in each image in the 561 nm channel by a local adaptive thresholding algorithm (Guanglei Xiong; Tsinghua University; http://www.mathworks.com/matlabcentral/fileexchange/8647-local-adaptive-thresholding). Subsequently the average distance of all peroxisome object pixels from a manually determined cell center was reported for each frame to monitor peroxisome movement over time. Additionally the change of the average 488 nm fluorescence intensity over the peroxisome object pixels was determine to measure motor recruitment to peroxisomes. Distance and intensity graphs were generated using the Origin software (OriginLab).

### In *vitro* microtubule gliding assay

TC-*Dm*KHC motors were expressed in bacteria, purified and then incubated (~10-20 μM) with 1 mM TCEP (tris(2-carboxyethyl)phosphine) for 5 min in elution buffer (50 mM NaPO_4_, 300 mM NaCl, 500 mM Imidazole, 1 mM MgCl_2_, pH 7.0) to reduce the thiol groups in the cysteines. Reduced motors were then incubated with approximately 5-fold excess TC-FlAsH II (catalog no T34561, Invitrogen) for 2 h at 4°C. Protein concentration and labeling stoichiometry was measured using a UV absorption spectrophotometer. Taxol-stabilized Cy5-labeled microtubules were adsorbed onto the surface of flow cells, and the surfaces were blocked with 2 mg/ml casein. Motility solution consisting of ~20 pM motors, 1 mM ATP, 0.2 mg/ml casein, 10 μM Taxol and an oxygen scavenging of 20 mM D-glucose, 0.02 mg/ml glucose oxidase, 0.008 mg/ml catalase and 0.5% v/v β-mercaptoethanol in BRB80 (80 mM PIPES, 1 mM MgCl2, 1 mM EGTA, pH 6.8) was then introduced. Microtubule gliding was visualized by TIRF using Nikon TE2000 microscope (60x, 1.45 NA PlanApo) equipped with a 488 nm Ar ion laser for GFP excitation and a 633 nm He-Ne laser for C5 excitation; experiments were performed at 26°C. Images were captured with a Cascade 512 CCD camera (Roper Scientific, Tucson, AZ) and acquisition and image analysis carried out using MetaVue software (Molecular Devices Corporation, Downingtown, PA); pixel size was 71.0 nm.

### Microtubule sliding assay in S2 cells

S2 cells were plated at 1x10^6^ per 1 ml media in 12 well dishes and co-transfected with tdEos-α-tubulin84B and BFP-tagged engineered kinesins (1 μg DNA total in a ratio of one to three). Immediately after transfection, 18 μg of double-stranded RNA targeting the 3’ untranslated region of KHC was added directly to the media. After 48 h, another 18 μg of dsRNA was added along with 200 μM copper sulfate to induce expression of the transfected constructs. Cells were imaged approximately 96 h after initial transfection and dsRNA treatment. On the day of imaging, transfected S2 cells were plated on Concanavalin-A treated coverslips 10 min before addition of either 0.3% EtOH/1.5 μM B-B Homodimerizer (Clontech) or 0.1% DMSO/400 nM ReAsH (Invitrogen) for 30 min. Cells treated with ReAsH were washed three times with 1 ml 1xBAL wash buffer in Insect-Xpress media. Before imaging of microtubule sliding, 2.5 μM Cytochalasin D and 20 nM Taxol were added to depolymerize F-actin and inhibit microtubule polymerization according to previous protocols^34, 48^. All imaging was performed within 30 min of Taxol addition. Photoconversion of a small subpopulation of microtubules was performed by 6 s exposure of 405 nm light from a Heliophor light source (89 North) constrained by a diaphragm. Photoconverted microtubules were imaged for at least 5 min with a 60 s interval using an inverted Nikon Eclipse U2000 microscope with a Yokogawa CSU10 spinning disk confocal head and Evolve EMCCD (Photometrics). Cells were photoconverted and imaged in groups of five using Nikon Elements software and a motorized stage. After photoconversion and imaging, BFP signal was imaged to confirm cells expressed wildtype or engineered kinesins. A custom Java-based Fiji plugin was developed to quantify microtubule sliding rates using the following methodology: Time-lapse images of photoconverted microtubules were bleach-corrected and thresholded to detect microtubules. The initial photoconverted zone was identified in the first frame of each movie. The number of pixels corresponding to microtubules was measured in total or outside the initial zone for each frame of time-lapse movies. The motile fraction of microtubules was calculated for each frame by: %MF = microtubules^outside_initial^_^zone^ / microtubules^total^. These values were then plotted against time and the slope of this graph was calculated for the initial linear section (identified by the highest R^2^ value of a linear regression which contained at least four data points). This slope represents the gross microtubule sliding rate in each cell with the units, Change in % Motile Fraction * min^−1^.

### *In vitro* single-molecule motility assays

Flow chambers (~10 μl volume) were constructed by attaching clean #1.5 cover glasses to microscope slides using double-sided adhesive tape. Microtubules were polymerized from a mixture of HiLyte647-labeled and unlabeled tubulins (Cytoskeleton) in BRB80 buffer (80 mM PIPES/KOH, pH 6.8, 1 mM MgCl_2_, 1 mM EGTA) in the presence of 1 mM GTP and 4 mM MgCl_2_ at 37°C and stabilized by the addition of 20 μM Taxol. Microtubules were diluted in five volumes of P12 buffer (12 mM PIPES/KOH, pH 6.8, 2 mM MgCl_2_, and 1 mM EGTA) supplemented with 10 μM Taxol and immobilized in flow chambers by non-specific adhesion. Flow chambers were subsequently incubated with 15 mg/ml bovine serum albumin in P12 buffer containing Taxol for 30-120 min at room temperature followed by motility mix containing 0.5-1.5 μl cell lysate in 30 μl of motility buffer (P12 buffer containing 17 μM Taxol, 1.7 mM MgCl_2_, and 1.7 mM dithiothreitol).

For each motor construct, motility assays in the absence and presence of inhibitor were carried out back-to-back. For ReAsH treatment, 0.5 μl of 2 mM ReAsH-EDT_2_ or 0.5 μl of DMSO was added to lysate in the motility mix and incubated 30-60 min on ice. Note that removal of serum albumin before making cell lysates is critical for inhibition as the dye binds non-specifically to BSA. For B/B treatment, 1.5 μM B/B homodimerizer or 0.5 μl ethanol was added to the lysate in the motility mix and incubated 30-60 min on ice. After incubation, 15 μl P12 buffer containing 30 mg/ml bovine serum albumin, 2 mM ATP and an oxygen scavenger system (final concentrations 0.08 mg/ml catalase, 10 mM D-(+)-glucose, 0.2 mg/ml glucose oxidase) was added to yield a final volume of 50 μl. The solution was then introduced to the flow chamber, and the chamber was sealed with molten paraffin wax.

Imaging was carried out at room temperature on a Nikon Ti-E/B microscope equipped with a 100x, 1.49 NA oil-immersion TIRF objective, 1.5x tube lens, three 20 mW diode lasers (488 nm, 561 nm, 640 nm) controlled via AOTF (Agilent, Santa Clara, CA), and EMCCD detector (iXon X3 DU897, Andor). Images were acquired with 488 nm excitation, 100 ms exposure at 9.83 frames per second. A still image of the microtubule track was acquired via 640 nm excitation before and after live imaging.

Single-molecule tracking^49^ yielded motor trajectories that were filtered based on temporal (minimum life time ≥ 0.5 s) and spatial (tracks fully overlap with the microtubule image) considerations. Motile events with a run length >250 nm were subjected to advanced trajectory analysis^50^. The run length was determined based on a start to endpoint vector, rather than a sum of the instantaneous positions along the trajectory. Such run lengths are insensitive to positional noise and are hence suited to analysis of run lengths of slow events e.g. for the inhibited TC constructs. Run lengths and run speeds of trajectories were plotted as empirical cumulative distribution functions and fitted to exponential (run length) and normal (run velocity) distributions to extract the means of these values. Ninety-five percent confidence intervals were obtained by bootstrapping (n boots = 2000). The motor landing rate was calculated as the number of motors observed to bind to the microtubule and processively moved for a distance greater than 250 nm normalized per μm of microtubule per minute.

## Miscellaneous

Structural Crystallography data were visualized with PyMOL (pymol.org).

## AUTHOR CONTRIBUTIONS

K.J.V. and W.O.H. conceived the inhibitable motor constructs, M.F.E, S.R., F.T., Y.Y., T.L.B., P.S. contributed to cloning, experiments and analysis of inhibitable rat KIF5C constructs *in vitro* and in cells, V.I.G. and M.W. contributed to experiments and analysis of *Dm*KHC constructs in S2 cells, W.O.H. and S.S. contributed to cloning, experiments and analysis of *Dm*KHC constructs in microtubule gliding assays. M.F.E and M.W custom coded analysis scripts. M.F.E. and K.J.V. wrote the manuscript with input from all authors.

## ACKNOWLEDGMENTS

We thank S. Norris and V. Soppina (Verhey laboratory) for support with TIRF imaging and data analysis; C. Hoogenraad (Utrecht University) and M. Rolls (Penn State University) for sharing plasmids. M.F.E. was supported by an early postdoc mobility fellowship (PBZHP3_141433) from the Swiss National Science Foundation. K.J.V. was supported by the NIH NINDS R21NS078761 and V.I.G. was supported by the grant GM-52112 from NIGMS.

## Supplementary Material

**Supplementary Video 1.** Microtubule-sliding in D. melanogaster S2 cells expressing tdEOS-tubulin. A subset of microtubules was photoconverted from green to red fluorescence in control (left) or KHC RNAi (right) treated cells. An image was captured in the red channel every min for 10 min. Scale bar, 5 μm.

**Supplementary Fig. 1.**
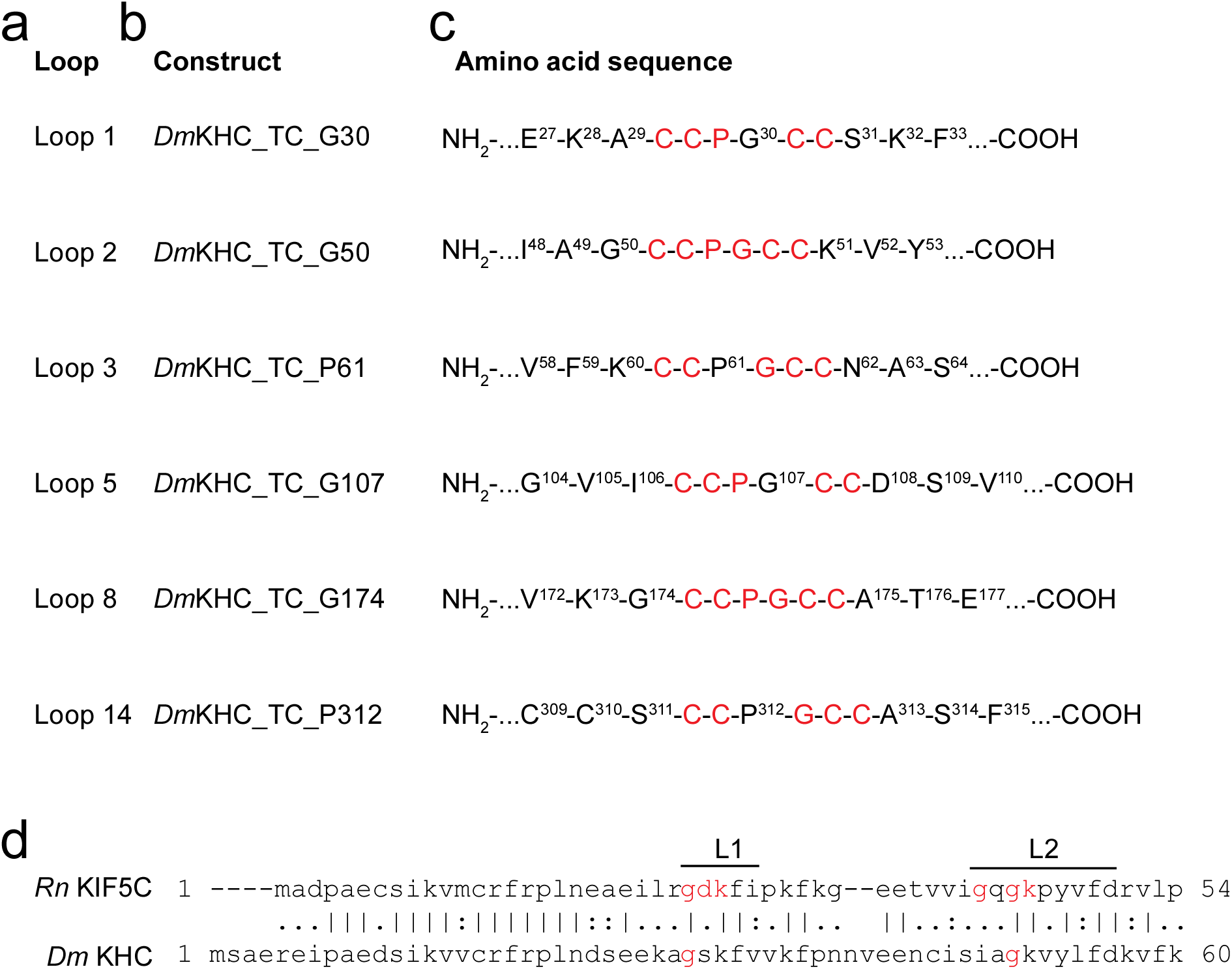
Design of TC-tagged *Dm*KHC constructs. (**a**) Surface loop location of TC insertion site. (**b**) Name of construct. (**c**) Sequence at TC (CCPGCC) insertion site (one letter amino acid code). Superscript numbers denote amino acid position in the protein; NH_2_, N-terminal; COOH; C-terminal. (**d**) Alignment of the N-terminal sequences of *Rn*KIF5C and *Dm*KHC using the Needleman-Wunsch algorithm (EMBL-EBI). A vertical line denotes identical, two dots highly similar, and one dot similar amino acids in the sequence comparison.

**Supplementary Fig. 2.**
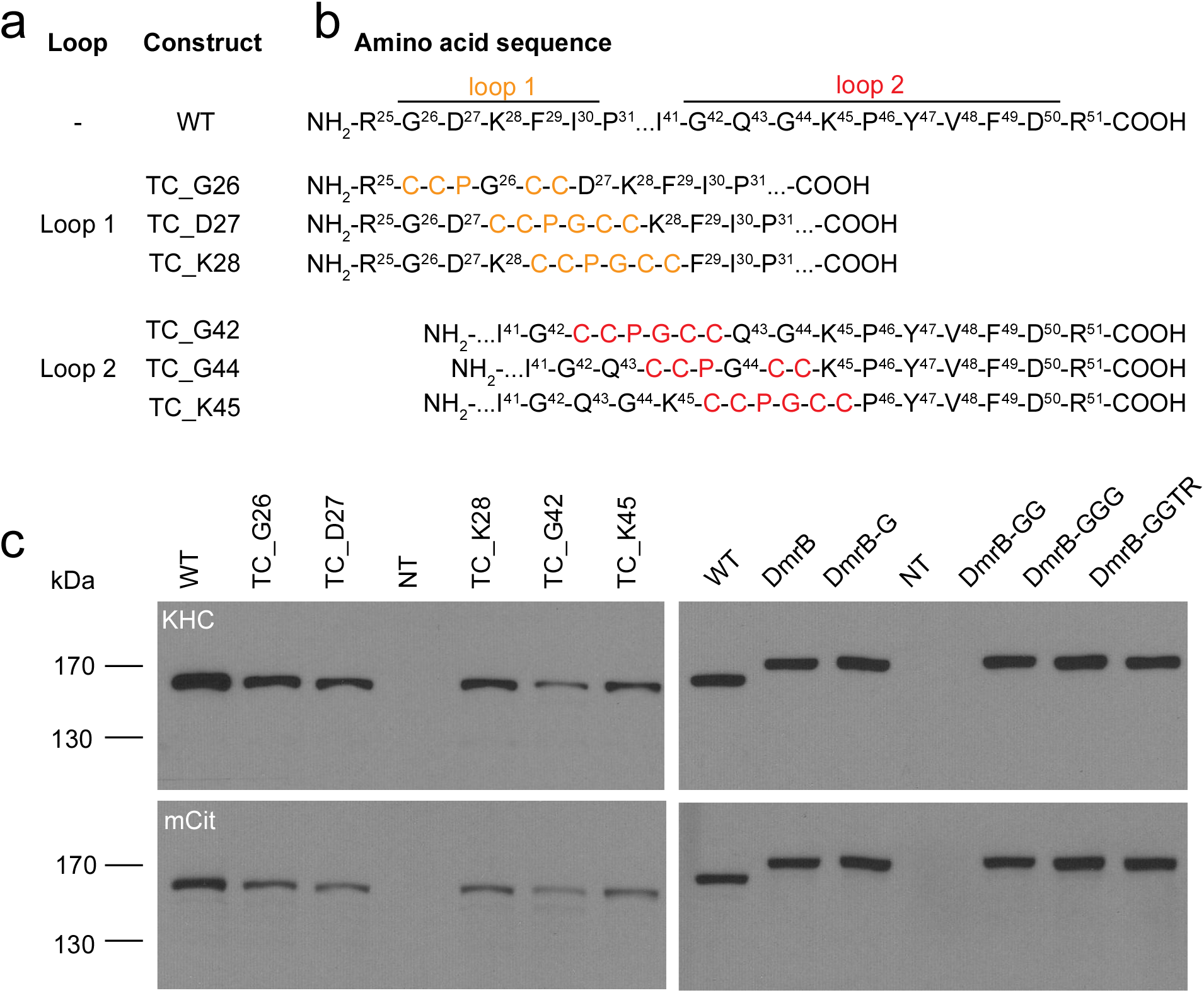
Design of inhibitable *Rn*KIF5C constructs. (**a**) Name of construct. (**b**) Sequence at TC insertion site (one letter amino acid code). Superscript numbers denote amino acid position in the protein; NH_2_, N-terminal; COOH; C-terminal. (**c**) Expression of inhibitable motor constructs in cells. Lysates of COS7 cells expressing the indicated TC-and DmrB-tagged constructs were immunoblotted with antibodies to kinesin-1 (H2; Covance MAB1614) and mCitrine (A6455; Invitrogen) as indicated. WT, wild type; NT, not transfected.

**Supplementary Fig. 3.**
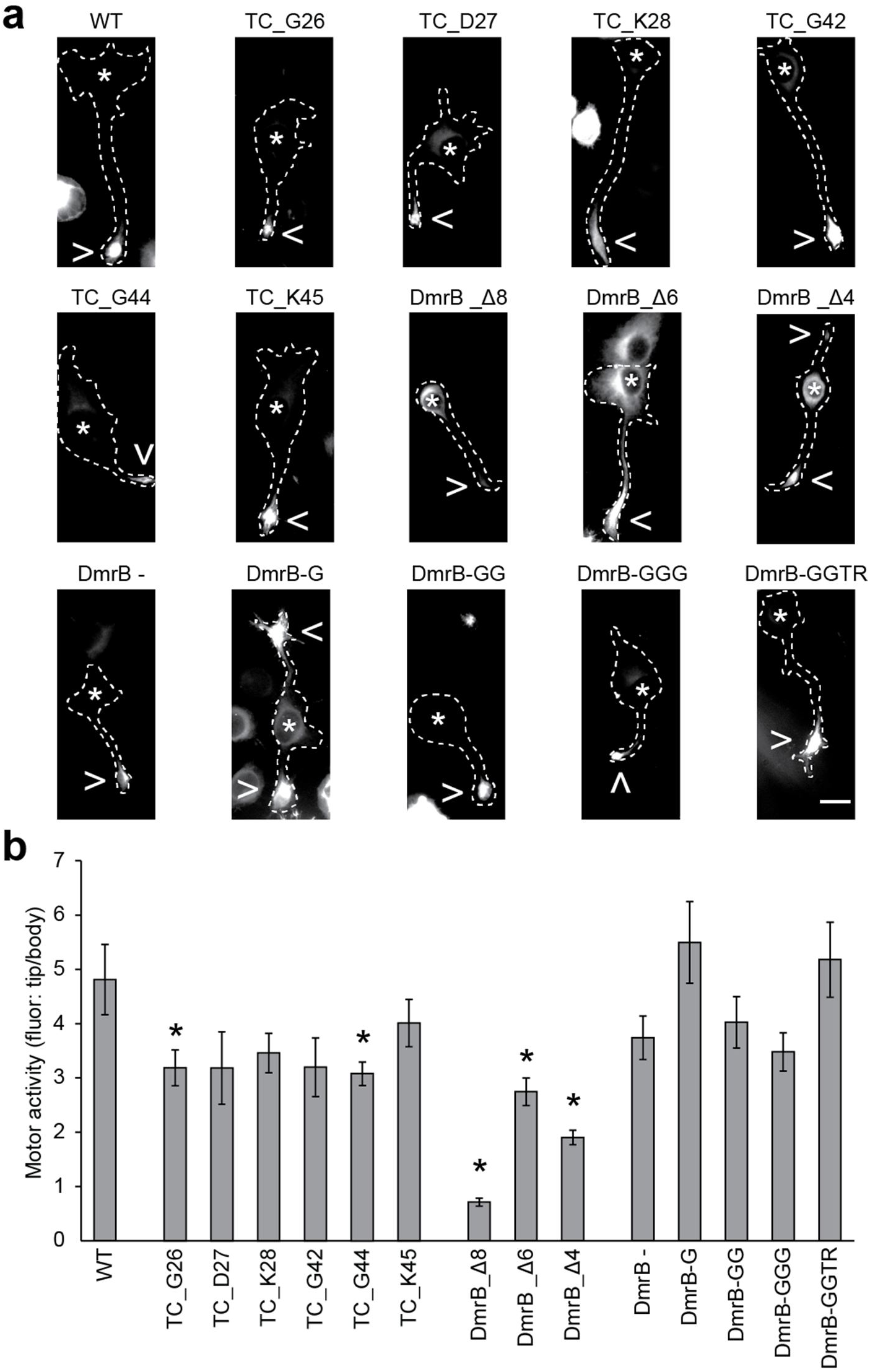
CAD cell tip accumulation assay. CAD cells transfected with mCit-tagged *Rn*KIF5C(1-559) motor constructs were serum starved to induce the formation of axon-like neurites. Two days later, cells were fixed and imaged. Scale bar; 15 μm. (**a**) Representative images showing the intracellular distribution of the indicated motor constructs. (**b**) Quantification of tip accumulation. The data are reported as the ratio of mean fluorescence in the neurite tip over the mean fluorescence near the cell body (n ≥ 19 cells each). A high ratio indicates an active *Rn*KIF5C(1-559) motor whereas a low ratio indicates an inactive motor. Error bars, SEM; * p<0.05 as compared to WT; p-values calculated with a heteroscedastic, two-tailed t test.

**Supplementary Fig. 4.**
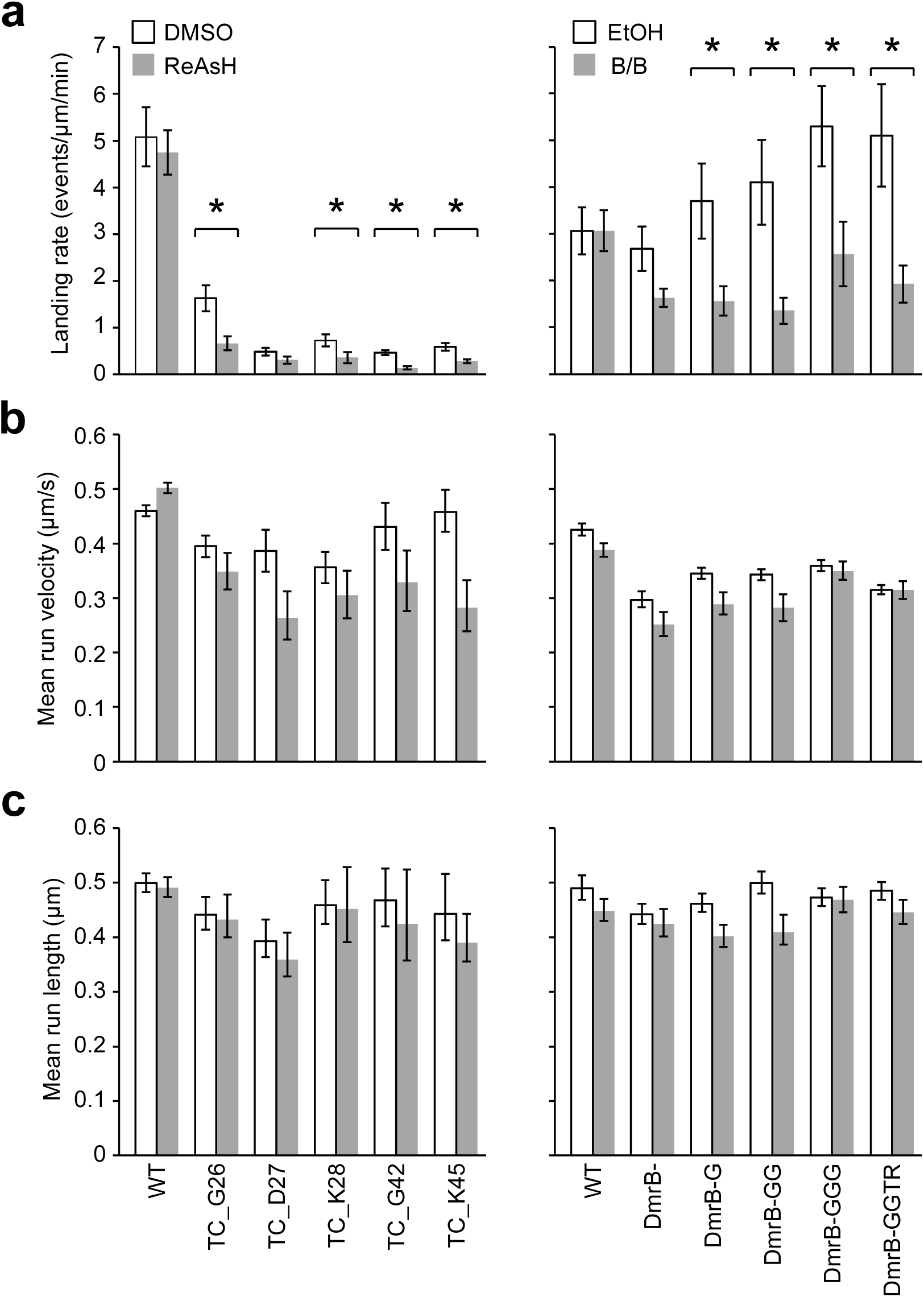
Single molecule motility properties of the engineered motors in the absence and presence of inhibitor. (**a**) The landing rate (number motility events > 250 nm) for each construct and treatment was obtained. Error bars, SEM; * p<0.05; heteroscedastic, two-tailed t test comparing motility in the absence and presence of drug. For the motility events, the mean (**b**) run velocity and (**c**) run length is shown. Error bars indicate 95% confidence interval obtained by bootstrapping.

**Supplementary Fig. 5.**
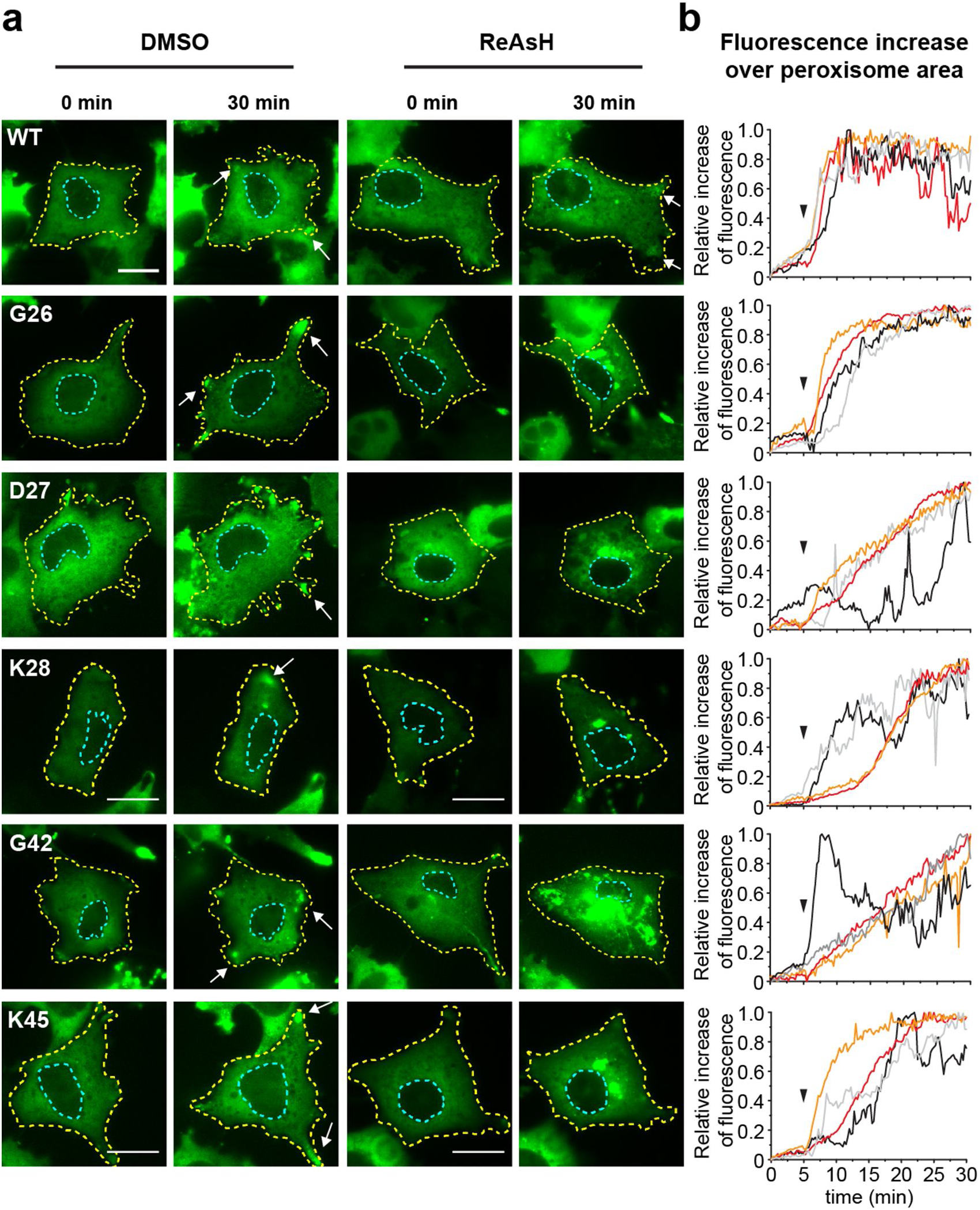
Motor recruitment to the peroxisome surface during the peroxisome redistribution assay. Cells expressing the indicated WT or TC-tagged motor constructs were treated with DMSO (vehicle control) or ReAsH for 30 min. Cells were then imaged for 30 min. Rapamycin was added 5 min after the start of imaging to induce motor recruitment to the peroxisome surface. (**a**) Representative images of the *Rn*KHC(1-559)-mCit-DmrC fluorescence at 0 min and at 30 min. Yellow dotted line, cell periphery; blue dotted line, nucleus. Arrows indicate the accumulation of *Rn*KIF5C-labeled peroxisomes in the periphery of the cell. Scale bar, 20 μm. (**b**) Time course of the relative increase in mCit fluorescence over the peroxisome area in the presence of DMSO (black and light grey lines) or ReAsH (red and orange lines). Rapamycin was added 5 min (arrow head) after the imaging start. The data are presented as the relative increase in mCit fluorescence over the peroxisome area normalized to the initial fluorescence and maximal fluorescence.

**Supplementary Fig. 6.**
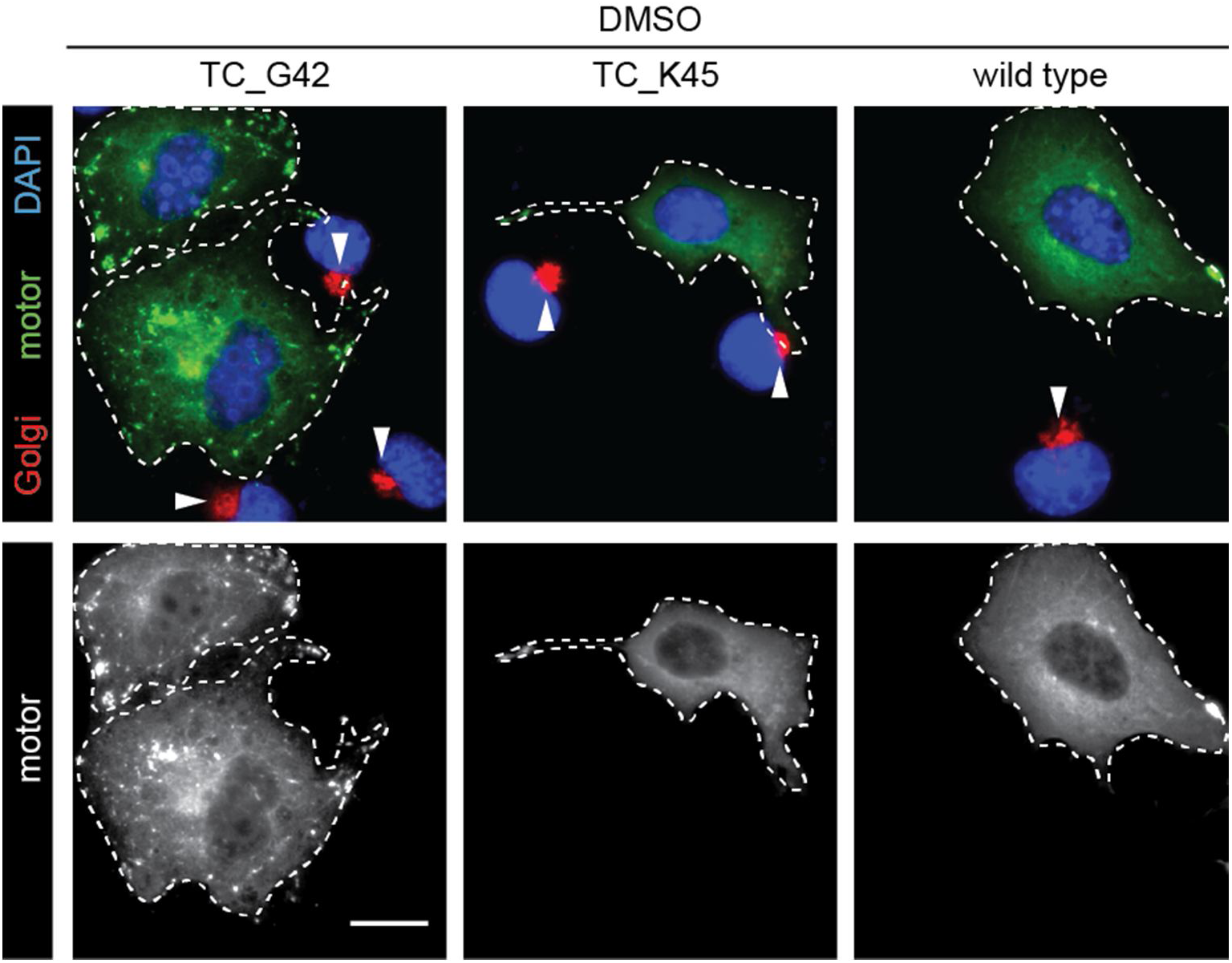
The TC_G42 version of *Rn*KIF5C(1-559)-mCit shows a tendency to aggregate in cells. Images of COS7 cells expressing WT, TC_K45 or TC_G42 versions of *Rn*KIF5C(1-559)-mCit-GTS (green) and fixed and stained with DAPI (blue) and an antibody against the Golgi marker protein Giantin (red). Top panels, merged images; bottom panels, motor (mCit) fluorescence. White dotted line, outline of cells expressing engineered motors. Arrowheads indicate compact Golgi complex in non-expressing cells. Scale bar, 15 μm.

**Supplementary Fig. 7.**
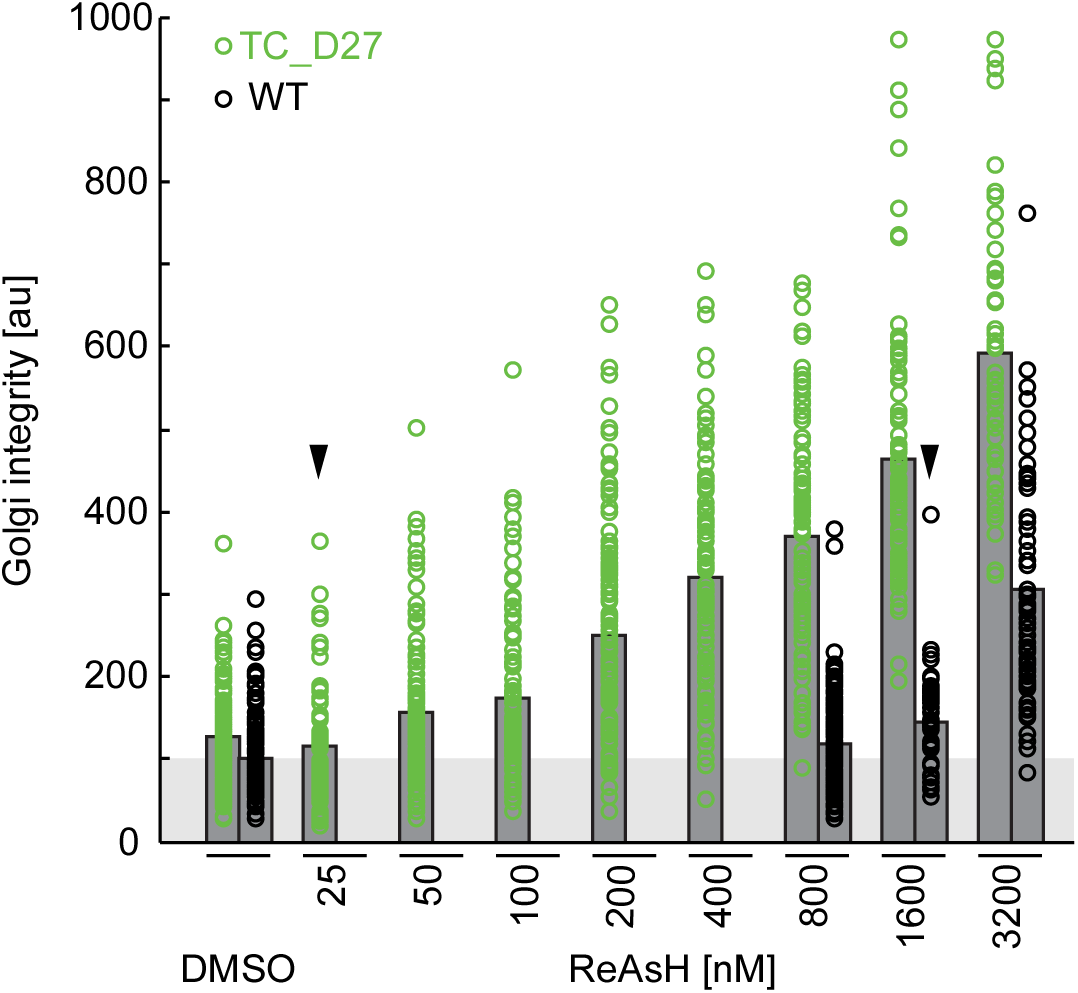
Dose-response curve for ReAsH inhibition in the Golgi redistribution assay. CO7 cells expressing WT or TC_D27 versions of *Rn*KIF5C(1-559)-mCit-GTS were treated with DMSO (vehicle control) or the indicated concentrations of ReAsH. 14 h later, the cells were fixed, stained with DAPI and an antibody to Giantin, and the Golgi integrity was measured. Low values reflect a dispersed Golgi complex (motors active) and high values reflect a compact perinuclear Golgi complex (motors inhibited). Data are from two independent experiments except when indicated by an arrowhead (one experiment). Each circle represents the Golgi integrity of an individual cell in WT expressing (black) or TC_D27 expressing (green) cells. Gray bars represent means.

